# SIPdb: A stable isotope probing database and analytical dashboard for linking amplicon sequences to microbial activity using a reverse ecology approach

**DOI:** 10.64898/2026.02.09.704843

**Authors:** Alex Batista Trentin, Abigayle Simpson, Jeffrey A. Kimbrel, Steven J. Blazewicz, Roland C. Wilhelm

## Abstract

Stable isotope probing (SIP) provides a powerful means to connect microbial sequence data with diverse metabolic activities, but the lack of a framework for SIP-derived data has limited its integration into broader strategies for ecological inference. Here, we introduce the SIPdb, an extensible SQLite database of curated nucleic acid SIP experiments (also in phyloseq format) paired with an interactive RShiny dashboard for analysis and visualization. The initial release compiles 22 studies covering 21 isotopolog substrates across diverse environments, with data standardized using the MISIP metadata standard. In creating the SIPdb, we have provided a standardized pipeline that accommodates the three most common SIP gradient fractionation strategies (binary, multi-fraction, and density-resolved), two isotope incorporator designation strategies (fixed- and sliding-window), and four complementary differential abundance methods (DESeq2, edgeR, limma-voom, and ALDEx2). Using our pipeline, we identified more than 42,000 unique amplicon sequence variants as isotope incorporators across 62 phyla. Benchmarking with synthetic datasets demonstrated consistent performance across incorporator designation strategies, with ALDEx2 providing the highest specificity. Validation against original publications showed that, on average, SIPdb recovered 70.1% of author-reported incorporator taxa, with discrepancies arising from differences in phylotyping or classification approaches. Finally, our reanalysis of a non-SIP study of 1,4-dioxane degradation showed how SIPdb can both validate known degraders and uncover additional candidate taxa involved in community metabolism. The SIPdb establishes a scalable platform for reverse ecology, enabling hypothesis generation, cross-study meta-analysis, and linking taxa to metabolic processes, while serving as an open, extensible resource to accelerate ecological interpretation in microbiome research.

## INTRODUCTION

Although the DNA sequencing revolution has significantly advanced our understanding of microbial diversity, efforts to interpret the ecological roles of community members remain limited by what can be called the ecological inference gap. This gap reflects a fundamental mismatch between commonly used sequencing methodologies and the inherent ecological and evolutionary properties of microbial systems, including variable gene copy numbers, strain-level heterogeneity, and the sparse availability of directly observed functions and activities.^1–4^ A major contributor to this gap is the reliance on culture-based characterizations as a proxy for ecological function, and the use of taxonomic identities to link to these established bodies of ecological knowledge. This approach is limited by the fact that many taxa detected through high-throughput sequencing lack experimentally verified ecological or functional traits, and by the ambiguity that results from reclassifications of taxonomic groups over time, including a recent revision of prokaryote nomenclature.^5^ Resources are needed that move beyond dependence on individual studies and literature review, allowing ecological interpretations and assumptions about microbiome patterns to be tested directly. In this context, reverse ecology offers a promising approach that centers on the relationships between genetic sequences, their environmental distributions, and their *in situ* functional associations.

Reverse ecology is an approach that infers the ecological roles and traits of microbes from patterns of gene or genome distribution across environmental gradients, host associations, or experimental conditions.^6,7^ Rather than focusing on characteristics of an organism, reverse ecology begins with a context, such as a soil disturbance or host, and identifies the genes, operons, regulatory elements, or genomes that are consistently associated with it. This strategy leverages the power of high-throughput sequencing not just to detect presence, but to identify environment-wide associations and ecological signals in large volumes of microbiome data. Recent studies have used reverse ecology to highlight the roles of rare microbes in adaptation to diverse abiotic factors^8^ and functional genes in relation to soil properties,^9^ and to associate specific bacterial taxa with responses to tillage and irrigation regimes in agricultural soils.^10^ This approach is particularly well suited to pattern detection methods such as machine learning^11^ and gains further power when paired with experimental strategies like stable isotope probing (SIP), which directly link sequence data to *in situ* metabolic activity. The effectiveness and scalability of reverse ecology is enhanced by the availability of curated, context-rich sequence databases,^12–15^ resources that may support training of predictive algorithms.

Nucleic acid SIP provides a powerful experimental complement to reverse ecology by directly linking specific microbial sequences with functional activities in natural contexts.^16,17^ In a SIP experiment, environmental samples (*e.g.*, soils, sediments, or hosts) are supplied with an isotopically labeled "heavy" substrate (*e.g.*, ¹³C, ¹⁵N, or ¹⁸O-enriched isotopologs). Organisms that metabolize the labeled substrates, or their by-products (i.e., ‘cross-feeding’), incorporate the heavy isotopes into their nucleic acids, which can then be separated based on their density shift and sequenced to designate ‘incorporators’ amplicon sequence variants (ASVs) or taxa.^18–22^ This technique circumvents the need for inferring culture-based or taxonomic traits and instead provides a direct link to metabolic activity, though caveats do apply (see Current Limitations section). SIP experiments have revealed the ecological roles of previously uncultivated microbes in processes such as degradation of complex organic matter,^23^ microbial growth and turnover,^24,25^ and trophic interactions like predation.^26,27^ The cross-study reuse has underscored the potential of aggregated SIP data to reveal broader ecological patterns.^27^ Our research demonstrates how a curated database of nucleic acid SIP sequence data (‘SIPdb’) can enhance the efficacy of a reverse ecology approach by linking microbial taxa to isotope incorporation patterns that reflect metabolic activity and ecological associations under environmentally relevant conditions.

Here, we describe the development and validation of the SIPdb, a comprehensive and extensible database designed to synthesize SIP-derived amplicon data for designating isotope incorporators across diverse substrate isotopolog experiments. The SIPdb accommodates three primary types of nucleic acid SIP gradient fractionation strategies: simple binary (‘heavy’ vs. ‘light’), position-resolved, and density-resolved; and two isotope incorporator designation strategies: fixed- and sliding-window. In curating the SIPdb, we adopted the Minimum Information for any Stable Isotope Probing Sequence (MISIP) standard,^28^ ensuring future SIP data can be easily integrated. While the current release is based on V4-region 16S rRNA gene data, the SIPdb framework can be extended to accommodate full-length amplicons and whole-metagenome SIP datasets, allowing for greater genomic resolution as more sequencing data become available. To accompany the SIPdb, we developed an interactive RShiny dashboard (www.sip-db.com) that enables researchers to explore, compare, and analyze their own microbiome data against curated SIP datasets. The dashboard supports users to perform sequence homology and taxonomy-based queries, along with data visualization. While SIPdb is not a quantitative SIP (qSIP) resource and does not provide absolute incorporation rates,^29^ it accommodates the most commonly generated nucleic acid SIP data types. Most importantly, it was designed to extend ecological interpretation to non-SIP microbiome datasets, offering a scalable and empirically grounded framework for inferring putative ecological associations and metabolic participation from SIP-derived sequence data, as demonstrated in our study.

## METHODS

### Obtaining and curating SIP amplicon sequencing datasets

The initial release of SIPdb integrates data from 22 SIP studies encompassing 21 amended isotopologs and 14 distinct environmental biomes (**Table S1**). In total, the database comprises 4,770 amplicon sequencing libraries and 163,417 unique ASVs, of which 46,484 were designated as significant isotope incorporators in one or more datasets (including H_2_^18^O) under the most relaxed criteria (log_2_FC > 1, *p*_adj_ ≤ 0.05). Incorporator criteria can be adjusted to implement more conservative threshold to filter SIPdb associations (as described below). Candidate datasets were identified through a combination of searches in the NCBI Short Read Archive (SRA), peer-reviewed literature, and a curated compilation provided by Simpson et al. (2023). Raw sequencing files were retrieved from the SRA using fasterq-dump from the SRA Toolkit^30^ and processed through a standardized QIIME2 workflow to generate phyloseq objects for downstream analysis (details in Supplementary Methods).

Due to incomplete study metadata (see ref ^28^), a total of 22 high-quality SIP datasets met all inclusion criteria and were retained in SIPdb. For inclusion in the SIPdb (current and future), all datasets must have the minimum set of metadata required by MISIP: isotope, isotopolog, isotopolog label, isotopolog approach, and gradient position. To ensure the initial instantiation of the SIP was exemplary, we added quality thresholds stipulating that all datasets contain: more than one gradient position, and comply with sequencing depth and sparsity quality thresholds (see next paragraph). Metadata from all included studies were standardized following the MISIP framework (available in **Table S2**), ensuring interoperability and facilitating cross-study integration.

At present, only amplicon data overlapping the full V4 region of the 16S rRNA gene was kept, bounded by the primer set 515F (5’-GTGYCAGCMGCCGCGGTAA-3’) and 806R (5’-GGACTACHVGGGTWTCTAAT-3’).^31^ Raw sequence data were processed in QIIME2 (v. 2024.10),^32^ with quality filtering, denoising, and chimera removal performed using the DADA2 plugin, producing ASVs. Taxonomic classification was assigned against the SILVA 138.2 database.^33^ The resulting feature table was filtered to remove sparse and low-abundance taxa. ASVs were retained if they: (i) had at least 3 total counts across all original study samples, (ii) were present in at least two study samples or 1% of samples per study, and (iii) exhibited ≤99% sparsity across study samples. Additionally, samples with fewer than 1,000 total reads were excluded from differential abundance testing, along with the paired controls. These adaptive thresholds balance retention of rare taxa that may become isotope incorporators with removal of potential sequencing artifacts, while accounting for varying sample sizes across the 22 studies. Sequences identified as chloroplasts or mitochondria were removed. A rooted phylogenetic tree was generated to support downstream diversity analyses using the QIIME2 q2-phylogeny workflow align-to-tree-mafft-fasttree. Finally, the ASV table, taxonomic assignments, representative sequences, sample metadata, and phylogenetic tree for each study were imported into R (v4.4.1)^34^ and formatted into a phyloseq object (v1.48.0).^35^

### Approaches to SIP incorporator designation

We developed a largely automated pipeline to standardize the identification of isotope incorporators from nucleic acid SIP datasets based on the differential abundance of sequences, either ASVs or taxa, in nucleic acid pools from isotopically enriched ("labeled") and natural abundance ("unlabeled") samples. The pipeline is coded in R and accepts phyloseq objects containing (1) an ASV count table, (2) corresponding taxonomic assignments, and (3) MISIP-formatted metadata including any other study factors. Importantly, the incorporator-designation pipeline does not use the phylogenetic tree or the representative ASV sequences; these elements are included in the phyloseq object only to support downstream analyses (e.g., UniFrac-based diversity metrics) and are not involved in determining incorporator status. Using SIP metadata and the count table, isotope incorporators are designated based on one of three gradient fraction cases and two window approaches for *in silico* combining of fractions. The pipeline outputs standardized results for each dataset, including: (1) taxon-level statistics for differential abundance (log_2_fold-change, p-values, adjusted p-values) generated by multiple methods (DESeq2,^36^ edgeR,^37^ limma-voom,^38^ and ALDEx2^39^); (2) corresponding metadata describing incorporator designation strategy (heavy fraction, sliding window, or substrate-specific comparisons), and (3) complete taxonomic and study-level provenance information. These results are compiled into a master table and summary reports that are subsequently imported into the SQLite SIPdb database (see next section). All code to reproduce the database is available on GitHub: https://github.com/alexbtrentin/sipdb-pipeline.

To adapt to variation in SIP gradient fractionation approaches, the pipeline accommodates three different gradient structures when designating isotope incorporators: (case 1) ‘binary,’ where fractions are pooled into heavy and light categories; (case 2) ‘position-resolved,’ with multiple sequential fractions but no buoyant density information; and (case 3) ‘density-resolved,’ where fractions include buoyant density data (**Figure 1A**). To flexibly identify incorporators, the pipeline integrates four established differential abundance methods (DESeq2, edgeR, limma-voom, and ALDEx2) and applies two complementary strategies for contrasting differential abundance between labeled and unlabeled controls to designate incorporators: a fixed-window and sliding-window approach, as previously described.^40^

**Figure 1.**
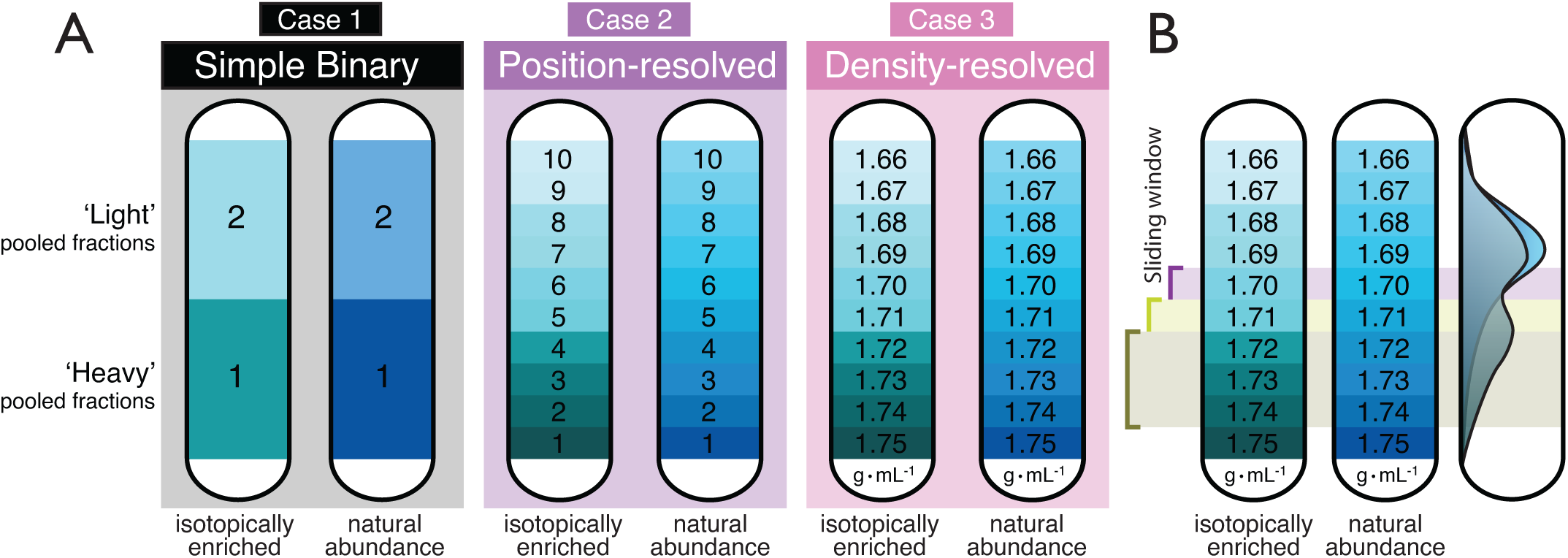
(A) Schematic representation of the three SIP study types supported by SIPdb: Case 1 (simple binary comparison of pooled “light” and “heavy” fractions), Case 2 (position-resolved gradients with discrete fraction numbers), and Case 3 (density-resolved gradients with measured buoyant densities). (B) Illustration of the sliding-window strategy applied to position- and density-resolved datasets, showing how overlapping windows capture local enrichment patterns while preserving gradient structure.

Pooling fractions (*in silico*) along the SIP gradient can improve sensitivity for detecting isotope incorporators, particularly when individual fractions have low sequencing depth. Pooling is also essential for case 2 studies, where density information is lacking and corresponding positions may differ slightly in buoyant density. By aggregating adjacent gradient positions, contrasts between labeled and unlabeled samples become less affected by stochastic variation and yield more reliable statistical inference. We implemented two window-based pooling strategies: a fixed-window approach, which designates the heaviest fraction range based on a consistent proportion of available positions within each study, and a sliding-window approach, which sequentially compares sets of adjacent gradient positions across the full gradient (**Figure 1B**).

To validate these approaches, we applied both fixed- and sliding-window methods to all case 2 (*n* = 14) and case 3 (*n* = 4) datasets and compared their relative information loss or gain. For position-resolved studies without density information (case 2), the fixed-window approach grouped the heaviest 33% of fractions from isotopically labeled samples as the “heavy” window and compared them directly against the corresponding heaviest 33% of fractions from the natural abundance controls, while lighter fractions from both labeled and unlabeled gradients were excluded from the analysis, following the rationale of Youngblut et al., 2014.^41^ When applying the sliding-window approach to position-resolved studies with few fractions (≤ 3), each fraction was analyzed as an individual window. For position-resolved gradients with ≥5 fractions, hierarchical clustering (Ward’s method with Euclidean distance)^40^ was used to group adjacent positions (e.g., 2–3–4) into two to five windows. These groupings were applied only to contiguous gradient positions, preserving the physical ordering of fractions along the gradient. An alternative k-nearest-neighbor (KNN)^42^ sampling approach was available for specific cases, adaptively selecting equal numbers of labeled and unlabeled samples (default: k=3 per group, adjusted based on total sample number) closest to each gradient position, maintaining statistical balance while capturing localized enrichment patterns. For density-resolved studies using the fixed-window method, windows were primarily focused on the heavy fraction (≥1.725 g/mL for stringent analysis, ≥1.700 g/mL for relaxed analysis), with optional borderline windows (1.710-1.725 g/mL) for intermediate incorporators. Sliding-window analyses for case 3 used the continuous density measurements directly and did not require clustering. Both windowing strategies were applied to all datasets to compare their performance and were included in the final version of the database.

Results from each window approach and each gradient case were compared within each gradient case to quantify how the choice of window strategy affected incorporator detection. For each taxon and method, effect sizes across windows were combined using inverse variance weighting [weights = 1/max × (*p*_adj_, 0.01)], and *p*-values were combined using Fisher’s method. Direction consistency was assessed as the proportion of windows showing the same direction of effect. Cross-method consensus was established by combining method-level meta-*p*-values using Stouffer’s Z-score method. Taxa were classified as incorporators if they exhibited consistent enrichment across windows (agreement score > 0.5), met statistical thresholds (*p*_adj_ <0.05), and showed substantial effect sizes (|log_2_FC| >1).

To determine which windowing strategy produced the most reliable incorporator calls and to create a unified, high-confidence output for SIPdb, we compared the fixed-window and sliding-window approaches within each gradient case. For every taxon, evidence across all windows was summarized using the four differential abundance methods implemented in our pipeline (DESeq2, edgeR, limma-voom, and ALDEx2). Window-specific effect sizes were combined using inverse-variance weighting, where weights were set to 1 divided by the maximum of the adjusted *p*-value and 0.01, and window-level *p*-values were combined using Fisher’s method.^43^ Direction consistency was calculated as the proportion of windows in which the taxon showed enrichment in the same direction, and we required a consistency score greater than 0.5 for inclusion. Outputs from the four statistical methods were integrated using Stouffer’s Z-score approach to generate a single cross-method consensus *p*-value for each ASV. Within each method, results from multiple density windows were first consolidated into a method-level log_2_-fold change (log_2_FC) estimate and a consistency score describing window behavior. The consensus log_2_FC reported in SIPdb corresponds to the mean of these method-level log_2_FC values, while the agreement score represents the mean of the per-method consistency scores. Results from multiple differential abundance methods and windowing approaches were thus integrated to produce a single consensus incorporator decision per ASV per study and substrate, while retaining full method- and window-level provenance in the database. Both fixed-window and sliding-window analyses were performed for case 2 and case 3 datasets, but only the sliding-window summaries were used to compute the consensus statistics incorporated into version 1 of SIPdb. Taxa were designated as final incorporators when they simultaneously met three criteria: a consensus adjusted *p*-value < 0.05, an absolute consensus log_2_FC > 1, and an agreement score > 0.5. When users select the ‘consensus’ option in SIPdb, these consensus statistics are displayed alongside all underlying window records for transparency, resulting in multiple rows per ASV that share the same consensus values rather than representing independent consensus calculations. Additional details are available in the Supplementary Methods.

### Validation of isotope incorporator designation with synthetic microbiome data

We evaluated the consistency, sensitivity, and accuracy of incorporator designation by generating synthetic microbiome datasets with defined compositions and processing them through the analysis pipeline. We generated 120 synthetic datasets spanning three gradient configurations: binary (n = 30 datasets), position-resolved with 3–8 discrete positions (n = 45), and density-resolved gradients (n = 45). Synthetic communities varied in richness (100–220 ASVs per dataset), sample number (16–48 samples), and isotope incorporation patterns. For each dataset, 5–18% of ASVs were designated as true incorporators, with simulated enrichment magnitudes of 2.8–5.5-fold in labeled samples from heavy fractions (buoyant density ≥1.725 g/mL or gradient positions 1–2). Biological noise was introduced through negative binomial count distributions with dispersion parameters typical of real amplicon data (size = 10). Natural abundance control samples were simulated without enrichment to mimic unlabeled experimental controls.

Each synthetic dataset was processed through our complete analysis pipeline (heavy fraction pooling, fixed-window, and KNN sliding-window analyses with all four differential abundance methods). Pipeline outputs were compared against the known ground truth data to calculate: (1) true positive rate (sensitivity/recall), the proportion of true incorporators correctly identified; (2) precision (positive predictive value), the proportion of predicted incorporators that were true; (3) specificity, the proportion of non-incorporators correctly excluded; and (4) F_1_ score, the harmonic mean of precision and recall, a balanced measure accounting for class imbalance. We systematically compared performance across fixed versus sliding-window approaches and the four differential abundance algorithms (DESeq2, edgeR, limma-voom, ALDEx2). Additional details are available in the Supplementary Methods.

### Validation against original SIP studies

To further evaluate the accuracy of SIPdb, we compared the designated incorporator results from our pipeline against those reported in the original study describing each dataset (n = 22 studies). For all included studies, we systematically reviewed the corresponding manuscripts and supplementary materials to extract the primary isotope-incorporating taxa reported by the authors. When available, ASV-level sequence identifiers were directly retrieved. Otherwise, genus- or family-level incorporators were compiled from tables, figures, or text descriptions. These author-reported incorporators were assembled into a curated reference list spanning 411 taxa across 21 isotopologs, which served as the basis for validation against SIPdb results (**Table S3**).

We then queried SIPdb to determine whether each reported ASV/taxon was detected as a significant incorporator (*p*_adj_ ≤ 0.05, log_2_FC > 1) using any of the four differential abundance methods (ALDEx2, DESeq2, edgeR, limma-voom), given the variability of approaches used among studies. A taxon was considered "validated" if it appeared in the SIPdb results for the corresponding isotopolog and study, regardless of which method(s) detected it (**Table S3**). For validated taxa, we recorded which methods identified the incorporator and the mean log_2_FC value. When available, ASV-level identifiers or representative sequences reported in the original publications were directly used for validation against SIPdb results. When ASV-level information was not provided, validation was performed at the taxonomic rank reported by the authors (typically genus or family). A reported taxon was considered validated if it appeared as a significant incorporator in SIPdb for the corresponding study and isotopolog using any differential abundance method. This mixed-resolution validation reflects reporting practices in the original studies and avoids excluding datasets lacking ASV-level identifiers.

### Constructing the SIPdb relational database

For the initial release of SIPdb, isotope incorporators were identified for each study dataset using the appropriate gradient case according to each study design. For position- and density-resolved studies, both fixed-window and sliding-window approaches were evaluated; however, only sliding-window consensus results were retained in SIPdb version 1 due to their greater stability across gradient configurations. Binary studies (case 1; n = 4) were analyzed as direct contrasts between heavy fractions of isotopically labeled and unlabeled samples. Position-based studies (case 2; n = 14) were analyzed using both fixed-window (pooling the heaviest 33% of fractions) and adaptive KNN sliding-window approaches. Density-resolved studies (case 3; n = 4) used the same dual windowing strategies, with windows defined by buoyant density ranges. All differential abundance analyses were performed on ASV counts, from which taxonomic incorporator designations were derived for extending cross-study comparability. An ASV was designated as an incorporator if it exhibited greater than 2-fold enrichment in the labeled nucleic acid pool (log_2_FC > 1) with Benjamini–Hochberg adjusted *p*-value (*p*_adj_) ≤ 0.05. We utilized dual density thresholds: the relaxed threshold (≥1.70 g/mL) to capture broader incorporation patterns with greater sensitivity, and a stringent threshold (≥1.725 g/mL) identifies high-confidence incorporators with reduced background. Both result sets are stored in the database (*sip results relaxed* and *sip results stringent* tables), allowing users to select thresholds appropriate for their research questions.

For each study, window, and substrate, differential abundance was evaluated using four complementary statistical frameworks (DESeq2, edgeR, limma-voom, and ALDEx2). All method- and window-specific results were retained in SIPdb with full provenance, including window definitions and sample composition. Results from multiple differential abundance methods and windowing approaches were integrated to generate a single consensus incorporator call per ASV per study and substrate, as described above.For SIPdb v.1, KNN sliding windows adaptively selected k = 3 samples per group (labeled and unlabeled) nearest to each gradient position. For sliding-window analyses, window metadata included position ranges (*window start*, *window end*, *window center*), window mode (KNN or fixed), and sample sizes per group. Studies were excluded if they had insufficient sample sizes (<2 per condition), extreme sparsity (>98% zeros), or lacked proper controls. Quality metrics captured for each study included the total number of samples per condition (*n labeled*, *n unlabeled*), median library size, data sparsity (% zeros), and the number of ASVs passing filtering thresholds.

Results were compiled into a relational SQLite database. The primary results tables (*sip results relaxed* and *sip results stringent*) link differential abundance results to study metadata (*studies* table), taxonomy (*taxonomy* table), and sequences (*asv_sequences* table) through foreign keys. Each results row documents what was tested (*ASV identifier*), how it was tested (*analytical method*), the comparison performed (*analysis type, comparison type*), gradient information (*gradient position, density threshold*), and window context for sliding analyses (*window name, window mode, k per group*). The *asv sequences* table contains all representative V4 region sequences with associated sequence length, GC content, and full taxonomic lineage.

To facilitate data access and analysis, SIPdb provides several curated tables: *analysis view relaxed* and *analysis view stringent* combine differential abundance results with study, substrate, and taxonomy metadata into a single queryable table; *enriched_sequences* lists all significantly enriched ASVs (*p*_adj_ ≤ 0.05 and log_2_FC > 1) together with their sequences and taxonomy; and *consensus_both_thresholds* identifies ASVs classified as incorporators under both relaxed and stringent density thresholds, providing a high-confidence subset for cross-study comparisons. All representative ASVs were compiled into a BLAST nucleotide database to enable homology-based queries using BLAST+ (‘all_sequences_v1.fa’).^30^ User-supplied 16S rRNA V4 sequences can be matched using standard BLASTn parameters, with incorporator status assigned based on sequence identity (≥97% identity for genus-level matches, or 100% identity over 251 bp for exact ASV-level matches).

Before finalizing version 1 of the SIPdb, we validated the following: (1) compared study-type detection accuracy (binary, multi-fraction, density classifications), (1) validated output completeness by confirming that the master table contained all required columns with expected data types and no missing critical values, (2) validated study coverage by verifying that all input datasets produced results in output files, and (3) validated file artifact integrity by checking that all expected output files (differential abundance tables, QC logs, summary reports, Excel workbooks) were generated without errors from all properly formatted phyloseq study inputs. Additional details are available in the Supplementary Methods.

### SIPdb analysis dashboard

To facilitate interactive exploration and broad accessibility, we developed a comprehensive Shiny web^44^ application that serves as the analysis dashboard for SIPdb (**Figure 2**), which is available to users at www.sip-db.com. The dashboard allows users to explore isotope incorporators across diverse isotopolog substrates and environmental contexts, and to annotate their own microbiome datasets using a web-based BLAST search against representative sequences of incorporator ASVs in SIPdb. Users can adjust statistical thresholds (*p*-value cutoff, log_2_FC ranges, relaxed vs. stringent density thresholds) and compare differential abundance methods (DESeq2, edgeR, limma-voom, ALDEx2). The SIPdb analysis workflow (phyloseq objects and R scripts), the SQLite database and sequence data, and the dashboard app (‘app.R’) are all open source and available for local use or adapting for future uses cases via GitHub.

**Figure 2.**
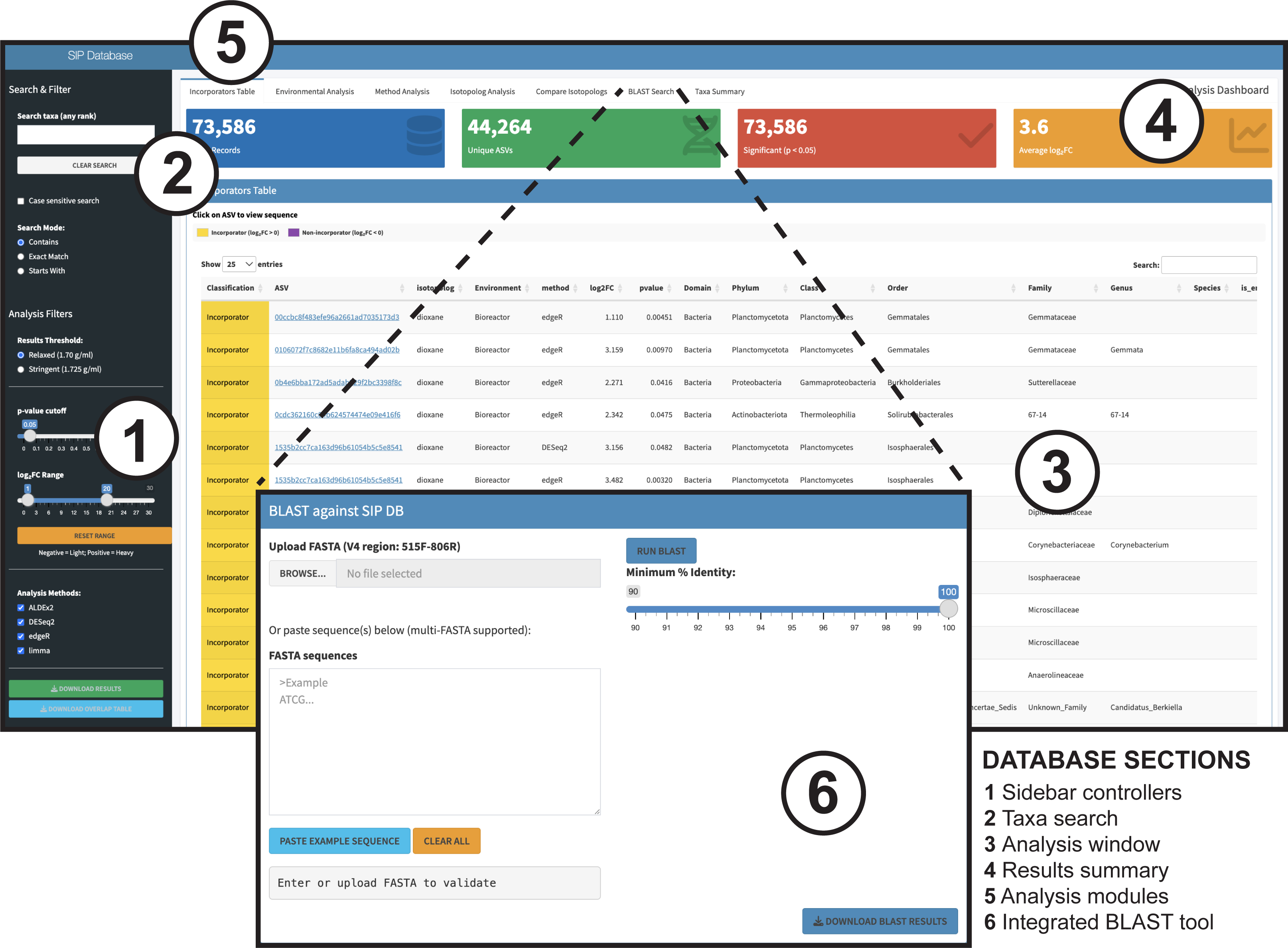
Overview of the SIPdb analysis dashboard. Screenshot of the interactive RShiny application developed for querying and visualizing SIPdb data. Key components include (1) sidebar controls for filtering by statistical threshold, isotopolog, and method; (2) search field for taxa or sequences; (3) results window displaying enrichment statistics and metadata; (4) summary panels showing study and dataset metrics; (5) module navigation bar for comparative analyses; and (6) integrated BLAST interface for matching user-provided 16S rRNA V4 sequences to SIPdb incorporators. The dashboard enables interactive exploration of isotope incorporation data and direct annotation of user microbiome datasets.

The dashboard comprises seven analysis modules. The *Incorporators Table* serves as the landing page, displaying ASVs meeting default enrichment criteria (*p*_adj_ < 0.05, log_2_FC > 1) with summary metrics (total incorporators, unique ASVs, studies represented), taxonomic search functionality, and exportable DOI-linked records. Five comparative analysis modules (*Environmental Analysis*, *Method Analysis*, *Isotopolog Analysis*, *Compare Isotopologs* and *Taxa Summary*) provide interactive visualizations including distribution bar charts with hover tooltips, summary statistics tables, and heatmaps of top 20 enriched taxa (by absolute log_2_FC), built with ggplot2,^45^ plotly,^46^ and viridisLite.^47^ The *Environmental Analysis* module enables users to examine how isotope incorporation patterns vary across environments, identifying environment-specific incorporator guilds. The *Method Analysis* module enables users to compare how different analytical pipelines (e.g., DESeq2, edgeR, ALDEx2) influence incorporator detection and effect size estimates. The *Isotopolog Analysis* module enables users to explore how different labeled substrates structure incorporator communities and enrichment strength across taxonomic ranks. The *Compare Isotopologs* module enables side-by-side comparison of taxon enrichment between any two substrates at user-selected phylotype or taxonomic ranks (ASV through phylum), with ranked bar charts and exportable overlap tables showing taxa present in both datasets. The *Taxa Summary* presents all records for a chosen organism at any taxonomic rank, with distribution plots across environments and isotopologs annotated with log_2_FC values.

Finally, the dashboard has a BLAST Search module which accepts user-provided 16S rRNA gene sequences in FASTA format (single or multi-FASTA), supporting up to 30,000 sequences or file sizes ≤30 Mb. Sequences must overlap the V4 region (515F-806R primers) to yield results. Queries are executed with BLASTn against the SIPdb sequence database, returning tabular alignments linked to incorporator metadata (study, method, log_2_FC, p-value, environment, isotopolog, full taxonomy). Results are filterable by sequence identity threshold (default ≥97%) and incorporator status (enriched in heavy vs. light fractions based on log_2_FC sign). All filtered tables and figures are exportable (CSV format), DOI fields link directly to primary publications, and ASV identifiers are clickable to display full V4 sequences. The application uses a Shiny/SQLite backend with planned regular database updates as new SIP studies become available. Additional details are available in the Supplementary Methods.

### Ecological Inference with SIPdb

SIPdb can be used in two primary ways to derive ecological insights: (i) conducting meta-analyses across SIP datasets, and (ii) mapping patterns of isotope incorporation onto non-SIP microbiome data to infer potential ecological associations or metabolic roles. To illustrate the first case, we visualized how substrate chemical properties influence microbial incorporation patterns across SIPdb by performing principal component analysis integrating substrate chemistry with incorporation metrics. For each substrate, we compiled six variables: (i) three chemical descriptors (solubility, hydrophobicity [xLogP3], and molecular weight from PubChem^48^, **Table S4**) and (ii) three incorporation metrics from significant enrichment results (*p*_adj_ < 0.05): number of incorporating taxa (unique ASVs/OTUs), mean differential abundance in isotopically-enriched DNA or ‘effect size’ (log_2_FC), and taxonomic breadth (number of unique phyla, a proxy for metabolic generality). Polymeric substrates (e.g., cellulose, DNA, chitin) were excluded because molecular descriptors are undefined for macromolecules with variable chain lengths, and substrates with incomplete chemical annotation were also excluded, yielding a total of 14 substrates in our analysis. Count-based variables were log_10_-transformed to address right-skewed distributions, and PCA was performed on scaled and centered data using prcomp in R^34^. To examine relationships between substrate chemistry and labeling strength, substrates were classified into effect-size categories (Low, Medium, High) based on tertiles of mean log_2_FC, and differences in PC1 scores among categories were assessed using Kruskal-Wallis tests with Dunn’s post-hoc comparisons (Benjamini-Hochberg correction).

To illustrate the second case, we evaluated whether SIPdb could aid interpretation of non-SIP microbiome datasets. We re-analyzed 16S rRNA gene data from Li et al. (2025),^49^ which examined two 1,4-dioxane–contaminated field sites with contrasting degradation outcomes (BioProject: PRJNA1183415). Data were processed using the same workflow applied to SIPdb studies. Briefly, raw reads were processed in QIIME2 (v2024.10) using DADA2 for denoising and chimera removal, yielding 13,176 ASVs, and taxonomy was assigned using SILVA 138.2.^33^ Putative 1,4-dioxane–responsive taxa were identified based on differential abundance (assessed using DESeq2, consistent with the analytical framework applied in the original study) between microcosms amended with 1,4-dioxane and unamended controls. Differentially abundant ASVs were queried against SIPdb using the integrated BLAST tool; however, few met the criterion of 100% identity across the full V4 region. In contrast, many ASVs that were not differentially abundant exhibited perfect sequence matches to SIPdb records. Consequently, a taxonomy-based matching approach was adopted. Differentially abundant ASVs from Li *et al.* were aggregated at the genus level, and genera were considered SIPdb-associated if any ASV classified to that genus had a matching record in SIPdb. Genera were then classified into three categories: *validated* (reported by Li et al. and present in the SIPdb match set), *recovered* (not reported by Li et al. but present in SIPdb as dioxane or phenanthrene incorporators), or *absent* (enriched in contaminated samples but not represented in SIPdb). SIPdb matches associated with other substrates (e.g., methane, butane, acetate) were retained for interpretive context only. Statistical comparisons used Wilcoxon tests for relative abundance and Fisher’s exact tests for presence–absence.

## RESULTS AND DISCUSSION

SIPdb represents a first step toward centralizing SIP-derived data within a unified database and analysis dashboard that facilitates cross-study comparison and supports ecological inference from standard amplicon-based microbiome datasets. All underlying methods, database structures, and analysis code are provided as open-source resources, enabling users to adapt, extend, and build upon the functionality implemented in version 1.

### Overview of SIPdb incorporator designations

Version 1 of the SIPdb currently contains 22 studies, representing a broad range of environments (14 different environment types) and substrates (21 isotope labelled substrates). The database identified 46,484 unique ASVs as significant isotope incorporators (*p*_adj_ ≤ 0.05, log_2_FC > 1). The isotopolog substrates span a broad range of metabolic contexts, including simple carbon substrates (glucose, methane), fermentation products (acetate, lactate, oxalate), complex carbohydrates (alginate, cellulose), aromatic compounds (phenanthrene, vanillin), and specialized substrates such as alkanes (butane), xenobiotics (dioxane, sulfamethoxazole), and biomolecules (DNA, amino acids). Within this set, ^18^O-labeled water occupies a distinct functional category, as it reflects the prevailing metabolic activity of all physiologically active organisms, independent of any specific substrate amendment. This allows ¹⁸O-H₂O to to serve as a baseline indicator of the actively growing microbiome within each environmental context, while underscoring the need to interpret SIPdb-derived incorporation patterns in light of the distinct ecological meanings associated with different isotopologs.

The SIPdb incorporator set spanned 62 phyla, with members of Pseudomonadota (27.6%), Bacteroidota (11.0%), and Chloroflexota (9.8%) representing the most frequently detected incorporators across studies. The predominance of Pseudomonadota as incorporators highlights the broad metabolic versatility and successful ecological strategies of this diverse lineage of bacteria, with members designated as incorporators across diverse substrates (**Figure 3**). Bacteroidota and Chloroflexota were also major isotope incorporators, though with a higher degree of substrate-specific associations.

**Figure 3.**
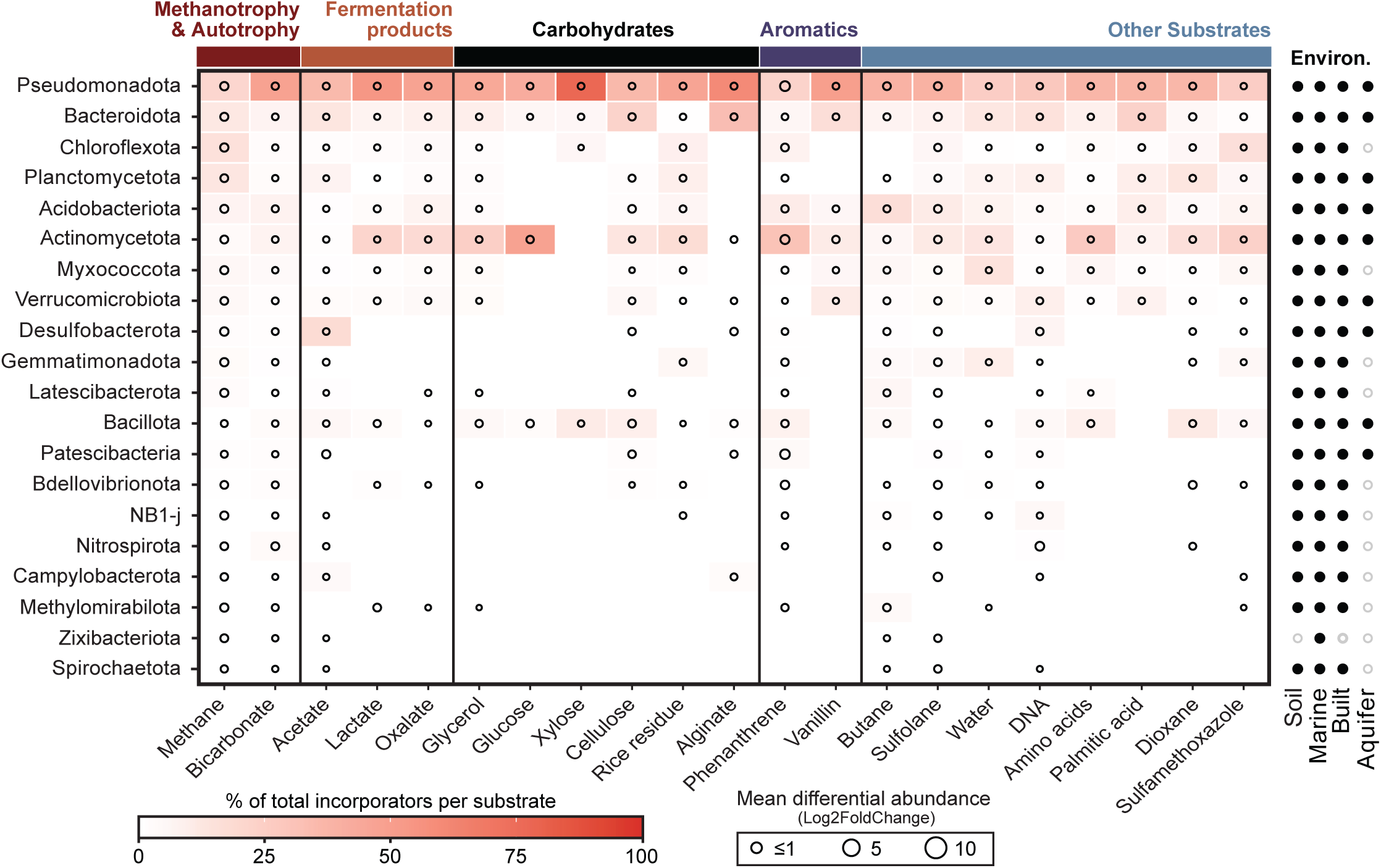
Cross-substrate distribution of isotope incorporators. Heatmap showing the relative abundance of isotope-incorporating phyla across 21 labeled substrates. Color intensity indicates the proportion of total incorporators per substrate, and circle size reflects mean enrichment (log₂FC). Substrates are grouped by chemical class, and the right panel shows the environments represented in each phylum.

Single-carbon (C₁) substrates yielded the highest numbers of designated incorporators, reflecting both the ubiquity of C₁ metabolism and the foundational role of these organisms in supporting carbon assimilation at higher trophic levels. Bicarbonate was incorporated by 3,633 ASVs across 41 phyla (mean log_2_FC = 2.35), indicating that the capacity to assimilate inorganic carbon is phylogenetically widespread, encompassing approximately 7% of ASVs detected in SIPdb. This proportion is higher than, but broadly comparable to, genome-based estimates suggesting that ∼2% of bacterial and archaeal genomes encode complete carbon fixation pathways.^50^ Importantly, our result is likely an overestimate given it likely reflects a combination of primary autotrophy, mixotrophy, and downstream incorporation via cross-feeding and trophic transfer, processes that cannot be disentangled using SIP alone. Consistent with this interpretation, Chloroflexota exhibited strong methane incorporation (3,624 ASVs), highlighting their potential importance within methanotrophic and methane-supported microbial communities.^51^ Many methane studies in SIPdb spanned long incubation periods (24 to 720 h), which likely promote secondary consumption and food-web propagation of ¹³C, explaining the large apparent taxonomic breadth of methane incorporators despite the taxonomically narrow set of known primary methanotrophs. In contrast to C1 compounds, a relatively narrow set of phyla were designated as glucose incorporators in SIPdb, unexpected given that glucose uptake is broadly distributed among bacteria.^52^ These observations suggest that growth traits (uptake kinetics and specific growth rates), substrate concentration, incubation time, and bioavailability, not just the presence of metabolic pathways, determine whether a taxon is designated as an incorporator.^48,53^

Together, the contrasting breadth of incorporators observed for C₁ substrates versus glucose highlights the ecological and methodological complexity inherent to SIP-derived incorporator data, emphasizing that incorporator breadth in SIP experiments reflects ecological context and trophic dynamics rather than simple substrate–pathway relationships.

### Benchmarking incorporator designation with synthetic SIP datasets

Across 120 synthetic datasets spanning binary (n = 30), density-resolved (n = 45), and position-resolved (n = 45) gradient cases, the pipeline performed well in distinguishing true incorporators from non-incorporators, with mean F_1_ scores comparable across gradient configurations: 0.784 for binary, 0.768 for position-resolved, and 0.815 for density-resolved gradients (**Figure 4A**). The three gradient types showed no significant performance differences (Kruskal-Walli’s test, *p* > 0.76 for all comparisons), indicating robust performance regardless of experimental design. Precision, sensitivity, and specificity were remarkably stable across gradient cases (**Figure 4B**), where precision (the proportion of predicted incorporators that were correct) remained consistently high at 0.889–0.904, sensitivity (the ability to detect true incorporators) ranged from 0.708–0.775, and specificity (the ability to avoid false positives) remained uniformly high at 0.983–0.988. These results demonstrate that the pipeline maintains balanced performance across different SIP experimental designs.

**Figure 4.**
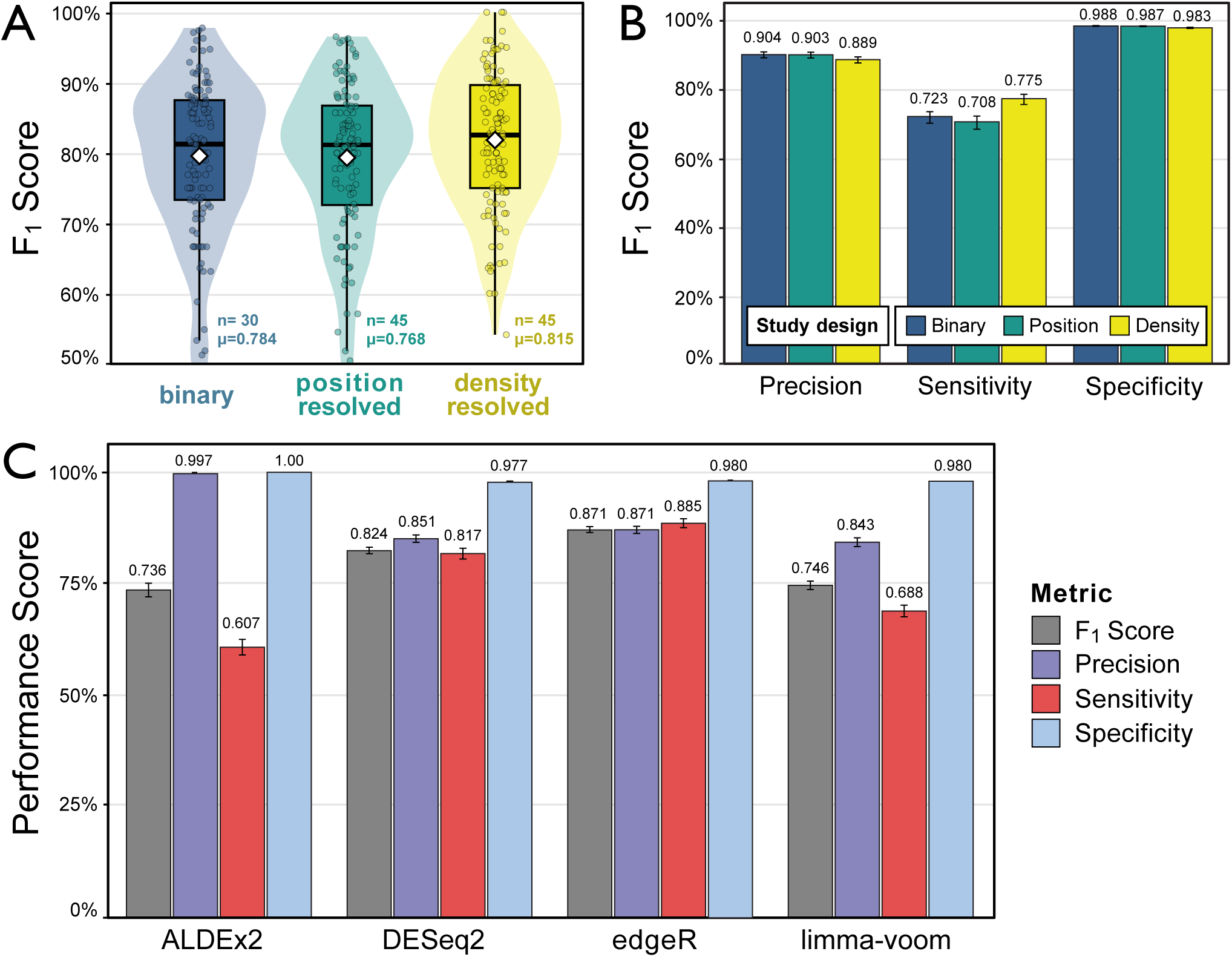
Overall performance of the pipeline in spiked mock data, with 30 generated mock objects per study type. A) Violin plot presenting the variation, and mean values, of the predictive performance (F_1_ score) among the three different study types. B) Different performance metrics for each of the three different study types. C) Overall predictive performance and metrics on each of the DA methods used on the database.

Accurate identification of incorporators varied substantially among differential abundance algorithms (**Figure 4C**), though all methods achieved relatively high precision (0.843–0.997) in this benchmarking. edgeR yielded the best overall F_1_ score (0.871) with balanced precision (0.871) and sensitivity (0.885), demonstrating strong all-around performance. DESeq2 achieved comparable performance (F_1_ = 0.824) with slightly higher precision (0.851) but lower sensitivity (0.817). ALDEx2, while exhibiting the highest precision (0.997) and specificity (1.000), showed the lowest sensitivity (0.607), reflecting highly conservative behavior that minimizes false positives at the cost of missing true incorporators, as previously shown ^54^. Limma-voom showed intermediate performance (F_1_ = 0.746) with moderate precision (0.843) and sensitivity (0.688).

edgeR’s superior F_1_ score, and its high level of sensitivity, in benchmarking is reflected in the large share of incorporator designations it contributes to the SIPdb (94% of all ASVs; 43,535/46,484). While this number is likely inflated, we configured SIPdb to display the union of all differential abundance methods, requiring users to explicitly select additional methods depending on the stringency desired. This design decision prioritizes identifying putative incorporators with a tolerance for false positives, a decision that should be revisited as more data is incorporated into the database. For the most stringent matching, we recommend using ALDEx2, which addresses compositionality with a robust statistical framework to handle source of false positives, though this conservatism results in lower sensitivity.^55^ The trade-offs among methods reflect fundamental statistical considerations: edgeR and DESeq2 maximize detection power, making them valuable when comprehensive incorporator identification is prioritized and downstream validation is feasible.^56^

These results indicate that our analysis pipeline is largely insensitive to study architecture (gradient type), with uniform performance across binary, density-resolved, and position-resolved configurations. The consistency across gradient types (mean F_1_ scores differing by only 0.047) underscores the robustness of the SIPdb analysis framework and its suitability for heterogeneous SIP study types. These findings also align with recent recommendations that differential abundance algorithm choice drives substantial variance in the performance of microbiome analyses.^57^

### Benchmarking against the original SIP studies

To validate SIPdb results against published findings, we systematically extracted the primary incorporators reported by authors in each of the 22 original studies, resulting in 411 taxa spanning genus to phylum level across 21 distinct isotopologs (**Table S3**). Of these author-identified incorporators, 70.1% (288/411 rank-agnostic-taxa) were successfully detected in SIPdb using our standardized analysis pipeline, demonstrating strong concordance between the database and original study conclusions. Detection rates varied substantially by substrate: perfect validation (100%) was achieved for alginate (8/8) and DNA (6/6), with high rates for water (88.9%), glucose (88.9%), sulfamethoxazole (88.9%), bicarbonate (85%), butane (83.3%), and methane (81.1%).

Lower detection rates were observed for palmitic acid (7.7%, 1/13) and xylose (8.3%, 1/12), both of which were reported in a single study in the current database. To determine whether these results reflected differences in statistical stringency rather than analytical failure, we performed a sensitivity analysis by relaxing the log_2_FC threshold to the modest > 0 while maintaining other criteria. Detection rates increased to 69.2% (9/13) for palmitic acid and 33.3% (4/12) for xylose, demonstrating that many author-reported incorporators produce detectable enrichment signals in SIPdb but with effect sizes below our conservative threshold.

Non-detection of author-reported incorporators (29.9%) likely stems from (1) statistical threshold differences between original studies and SIPdb’s standardized *p*_adj_ ≤ 0.05 criterion, (2) taxonomic assignment variation across reference databases and algorithms, (3) data processing heterogeneity in normalization and quality filtering, and (4) biological factors including stochastic sampling of low-abundance taxa. The strong overall concordance (70.1%) demonstrates that SIPdb successfully reproduces most published SIP findings while maintaining conservative statistical rigor, supporting its utility as a reliable resource for comparative SIP analysis.

### Cross-study exploration of substrate-association incorporator profiles

Principal component analysis revealed that substrate-associated incorporator profiles diverge according to substrate physicochemical properties and patterns of isotope incorporation breadth and strength (**Figure 5**). The first principal component (PC1) explained a large proportion of the variance (54.5%) and exhibited strong loadings for the proportion of incorporator taxa (0.502), the total number of incorporators (0.500), and taxonomic breadth (0.479). Together, these metrics capture variation in trophic accessibility, reflecting the extent to which a substrate’s isotopic label propagates through the microbial food web, from primary consumers to downstream cross-feeders^58,59^. Substrate positions along PC1 corresponded to significant differences in labeling strength (approximated by log_2_FC in relative abundance), where high-effect size substrates clustered at the high-trophic-accessibility end of PC1 (mean PC1 = 1.84), whereas medium-effect substrates occupied the low-trophic-accessibility region (mean PC1 = −1.73; Kruskal–Wallis χ² = 8.73, *p* = 0.013). The second principal component also explained a large proportion of variation (26.5%) with loadings associated with physicochemical accessibility, separating small, highly soluble substrates (e.g., acetate, glycerol) from hydrophobic or high-molecular-weight compounds (e.g., palmitic acid, sulfamethoxazole). Together, these axes demonstrate that substrate identity structures microbial communities not only through chemical constraints on uptake, but also through the extent to which isotopic label moves through trophic networks after primary assimilation.^60–63^

**Figure 5.**
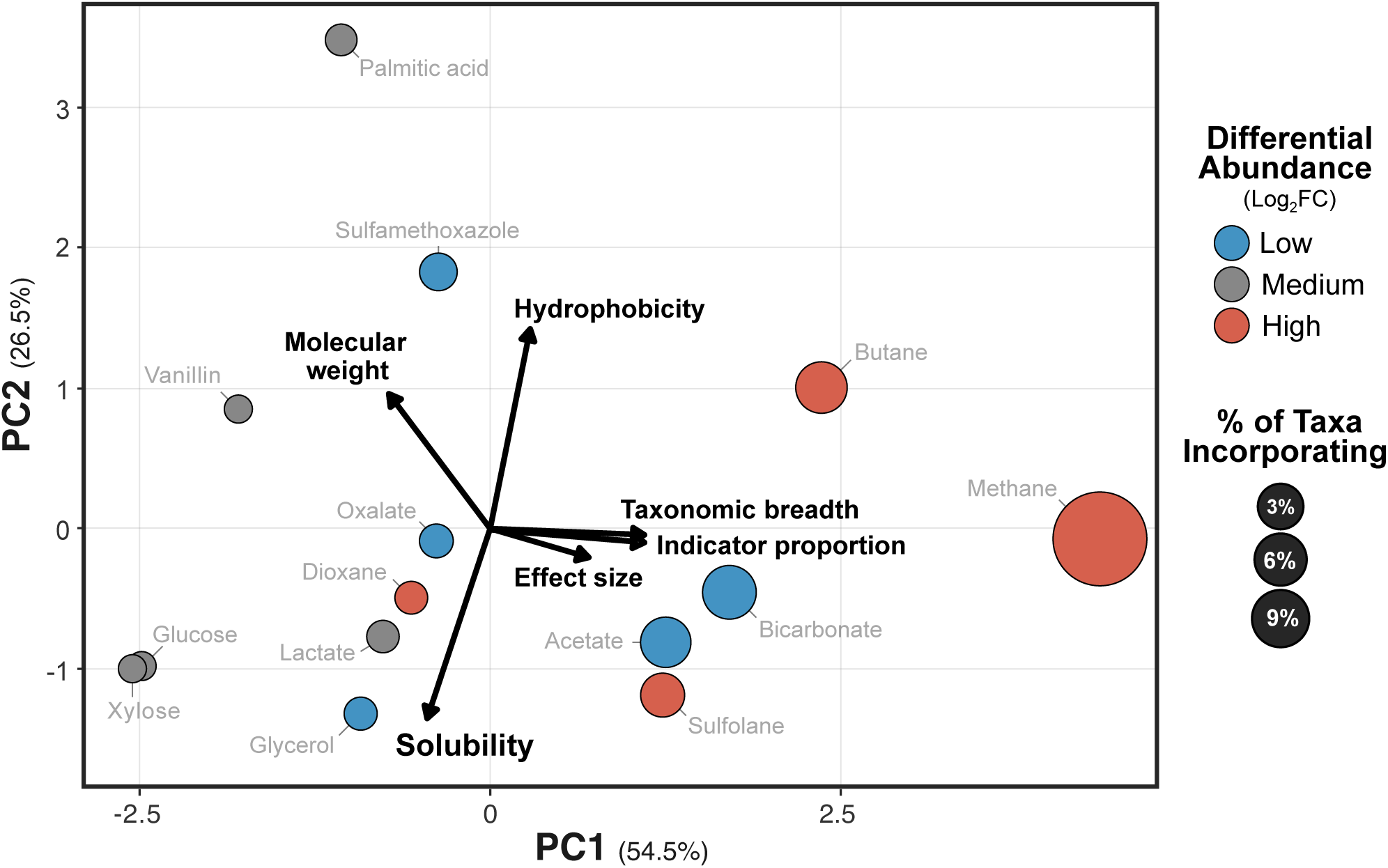
Principal component analysis of substrate properties and incorporator patterns. Biplot showing PCA of substrate physicochemical traits (e.g., solubility, molecular weight, hydrophobicity) overlaid with substrate-specific incorporator characteristics. Each point represents a labeled substrate, colored by mean effect size (log₂FC) and scaled by the proportion of ASVs designated as incorporators for that substrate relative to the total number of ASVs detected in the corresponding study or studies. Vectors indicate key physicochemical variables driving variation among substrates.

To evaluate reproducibility across independent studies, we examined substrates tested multiple times. Mantel tests revealed a significant positive correlation between study replication and phylum-level similarity (Spearman *r* = 0.256, *p* = 0.026), indicating that substrates examined across diverse environments, including methane and cellulose, yielded more consistent incorporator communities than those from single studies. This pattern was not driven by differences in within-substrate variance (betadisper *p* = 0.729). Reproducibility was strongly scale-dependent, with phylum-level profiles exhibiting substantially higher overlap across studies (mean Jaccard similarity = 0.44 ± 0.21) than genus-level profiles (0.039 ± 0.05), an 11.4-fold difference. This scale dependence mirrors broader patterns in microbial ecology, in which high-level functional traits are conserved within major clades, while fine-scale taxonomic composition reflects local environmental filtering^64–66^. Practically, this suggests that SIPdb information is propagated across broad taxonomic ranks, with phylum- and class-level guilds providing more reliable guidance for identifying likely incorporators of novel substrates than genus-level assignments. Together, these results show that SIPdb captures generalizable ecological rules of substrate use that extend beyond primary metabolism to include trophic accessibility.

### Reverse ecology approach using SIPdb

To evaluate whether SIPdb can inform mechanistic interpretation of non-SIP microbiome datasets, we mapped 16S rRNA gene profiles from a study of 1,4-dioxane degradation (Li *et al.*, 2025)^49^ onto SIPdb and assessed whether enriched taxa had prior isotope-based evidence of substrate assimilation. SIPdb currently contains one dioxane-degradation SIP dataset, conducted in activated sludge bioreactors, whereas Li *et al.* examined degradation in polluted groundwater sediments. Our goal was to assess whether SIPdb could help identify patterns in isotope-incorporating taxa, revealing genera with documented incorporation capacity that might otherwise be overlooked in site-specific analyses. Importantly, any matches we observe are likely to reflect both primary dioxane-degrading populations as well as cross-feeding taxa whose incorporation is driven by trophic propagation.

Exact OTU-level correspondence between datasets was rare, so a taxonomy-based approach was used to designate SIP incorporators. Of the 12 genera reported by Li *et al.* as enriched at dioxane-contaminated sites, five matched SIPdb records indicating incorporation by dioxane or the structurally related compound phenanthrene (**Figure 6**). These validated taxa were not randomly distributed within the community; they included several of the most abundant populations, such as *Brevundimonas* (26.1% of total reads, log_2_FC = 4.1), *Rhodococcus* (25.1%, log_2_FC = 2.7), and *Paenarthrobacter* (log_2_FC = 7.2; 9.8%). This pattern suggests that even with limited dioxane-specific data, SIPdb can help identify ecologically dominant degraders relevant to bioremediation applications. The validation of *Brevundimonas* and *Streptococcus* through direct dioxane SIPdb matches, alongside *Rhodococcus, Paenarthrobacter,* and *Lysobacter* through phenanthrene records, demonstrates the value of warehousing structurally related compounds in the SIPdb as means for discovery. The inclusion of phenanthrene-incorporating taxa in our query reflects the observation that bacteria capable of PAH degradation often possess diverse catabolic capabilities and broad-substrate monooxygenase systems.^67–69^

**Figure 6.**
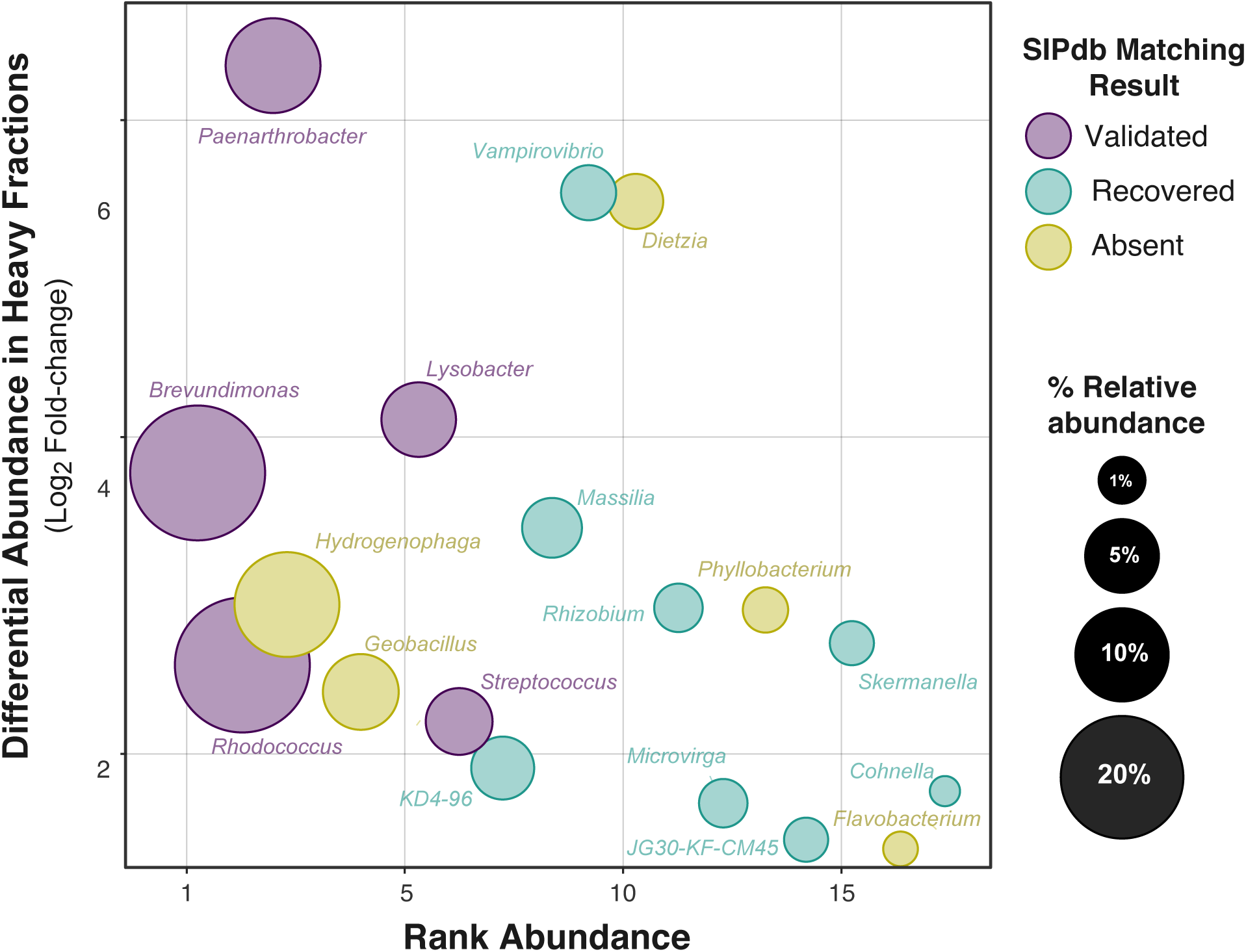
Reverse-ecology identification of taxa enriched at contaminated sites. Bubble chart showing taxa enriched in the 1,4-dioxane dataset, plotted by abundance rank (x-axis) and enrichment strength (log₂FC; y-axis). Circle size represents relative abundance, and color indicates discovery type: SIPdb-validated (present in literature and detected in SIPdb), SIPdb-recovered (not highlighted by authors but detected by SIPdb), and SIPdb-absent (reported by authors but not detected by SIPdb).

Five differentially abundant genera in Li *et al.* dioxane-amended microcosms were absent from SIPdb’s dioxane/phenanthrene records: *Dietzia, Hydrogenophaga, Phyllobacterium, Geobacillus,* and *Flavobacterium*. These taxa played an important role in the study system, with *Dietzia* exhibiting the second-highest relative abundance increase in their experiment (log_2_FC = 5.7), while *Hydrogenophaga* was among the most abundant (13.6% of total reads). Notably, three of the absent genera, *Hydrogenophaga, Flavobacterium*, and *Geobacillus*, were identified as incorporators for methane or butane incorporation in the SIPdb. This observation has important mechanistic implications: soluble di-iron monooxygenases, including propane monooxygenases and butane monooxygenases, exhibit broad substrate specificity and are well-documented for the cometabolic oxidation of dioxane.^70,71^ Taxa capable of C2–C4 alkane oxidation may therefore possess latent dioxane degradation capacity through these promiscuous enzyme systems.^72^ This result demonstrates how SIPdb may extend interpretation of results, enabling hypothesis generation beyond simple substrate matching. Querying taxa enriched for related pathways (e.g., C₂–C₄ alkane oxidation) allows users to infer co-metabolic potential even when direct SIP evidence is unavailable. Regardless, users may anticipate that phylotypes and taxa key to their study system may be absent from SIPdb due to either gaps in database coverage or geographical and ecological differences.Beyond confirming known degraders, SIPdb matches identified eight genera with documented dioxane and phenanthrene incorporation that were enriched at other contaminated sites but not reported in Li *et al*. Recovered genera included *Vampirovibrio* (log_2_FC = 5.7), Massilia (log_2_FC = 3.5), and multiple *Rhizobiales* members, including *Rhizobium, Skermanella*, and *Microvirga*. The presence of multiple Rhizobiales is notable given this order’s known metabolic versatility.^73,74^ *Vampirovibrio*, a predatory bacterium, may represent indirect enrichment through predation on primary degraders rather than direct dioxane metabolism. This fact illustrates the importance of interpreting matches to the SIPdb with caution, as the SIPdb can reveal both direct incorporators and taxa responding indirectly to contaminant exposure.^27,75^

Our reverse ecology approach demonstrates the use of SIPdb to drive hypothesis generation. By linking non-SIP observations to documented isotope incorporation patterns, SIPdb enables researchers to move beyond descriptive community analyses toward mechanistic inference. As the database expands with additional SIP studies across diverse substrates and environments, its capacity to inform and guide experimental priorities will continue to grow.

### Future advances and the promise of the SIPdb

Despite the relatively limited number of publicly available datasets in the SIPdb, due to legacy issues in data curation,^28^ our experimentation with the initial release of SIPdb revealed patterns in microbial substrate incorporation and demonstrated its promise as a tool for reverse ecology. All SIPdb entries are formatted according to the MISIP standard^28^, ensuring its forward compatibility with newly generated SIP datasets. MISIP is slated for adoption as a formal MIxS checklist (in version 7), which will guide SIP data archival for integration in the SIPdb. Furthermore, the volume of SIP sequencing data is expected to grow as recent innovations in high-throughput, semi-automated workflows reduce the labor burden associated with SIP, cutting hands-on time by six-fold.^60^ Previous advances in SIP computational methods were integral to the development of the SIPdb (e.g., HTS-SIP^41^) and continued progress in this space, including the use of synthetic DNA internal standards for improved quantification accuracy,^76^ will further enhance the aims and utility of SIPdb. Together, these advances pave the way for fully automated SIP analysis workflows, built on common standards with consistent data structures. As a fully open-source project, SIPdb is designed to support and grow with this evolving ecosystem, serving as a scalable platform for meta-analysis, cross-study synthesis, and hypothesis generation in microbial ecology.

### Current limitations and methodological evolution

While SIPdb represents a meaningful step forward in enabling cross-study synthesis and reverse ecology using SIP-derived sequence data, several limitations of its current form should be acknowledged. First, the database is currently restricted to V4 region amplicon data which has limited phylogenetic resolving power. This limits its utility for identifying fine-scale functional variation, such as strain-level population dynamics or accessory gene content, which are increasingly recognized as important drivers of microbial ecology and biogeochemical function.^77–79^ Expanding the SIPdb to include shotgun metagenomic and metatranscriptomic datasets will be essential for improving both taxonomic resolution and direct functional inference. However, such data were still relatively scarce at the time of publication.

The SQLite database and RShiny dashboard were designed with modularity and extensibility in mind, allowing users to apply different differential abundance frameworks and customize enrichment detection strategies, such as qSIP.^29^ As the SIPdb grows, it will need to accommodate more complex sequence types, including longer amplicons or metagenomic data. The current BLAST-based search architecture may need to evolve toward more scalable alignment and indexing tools, as well as more platform with greater computational resources like Galaxy^80^ or KBase.^81^ To extend the utility of SIPdb, one promising direction is the integration of *in silico* SIP simulations. Tools like *MetaSIPSim*^82^ already offer the ability to generate realistic synthetic SIP metagenomic datasets, which could be embedded in the dashboard to test methodological hypotheses or train machine learning classifiers for incorporator detection. Likewise, simulation-based modules could support researchers in experimental design, helping to evaluate tradeoffs in incubation length, substrate concentration, or gradient resolution, similar to experimental efforts in optimizing qSIP protocols.^83^

An additional limitation lies in the abstraction of study-specific methodological detail into simplified categorical metadata fields used in designating the isotopic incorporators that form the basis of the SIPdb. While necessary for interoperability, this standardization introduces a degree of data loss, particularly around incubation conditions, substrate labeling strategies, and gradient fractionation schemes. These factors can influence the identity and detectability of incorporators, potentially biasing results presented in the SIPdb toward taxa exhibiting the strongest incorporation effects. This tradeoff between standardization and preserving useful contextual information reflects a broader challenge in microbiome database development, where ease of use and broad applicability must be balanced against preserving nuanced experimental context. Future iterations of SIPdb could include layered metadata or filterable provenance flags that allow expert users to drill down into methodological differences while retaining an accessible default view for simple exploratory analyses. These opportunities will grow with more explicit and guided metadata collection, as being advocated by the National Microbiome Data Collaborative (NMDC),^84^ as well as possibly AI-enhanced analyses. At the risk of overstating the value of additional data, the utility of SIPdb for cross-study meta-analysis and reverse-ecology applications is expected to expand as the database grows, enabling identification of recurring activity patterns associated with specific taxa or phylotypes and comparison of incorporation dynamics across diverse environmental contexts.

## ACKNOWLEDGEMENTS

We thank Kaitlyn Johnson and Payton Taylor, at Purdue University, for their help curating the list of SIP studies. We would like to thank the Brazilian Ministry of Education and the Coordenação de Aperfeiçoamento de Pessoal de Nível Superior (CAPES) Fellowship Program (https://ror.org/00x0ma614) for the generous funding support for ABT. We would also like to thank the USDA National Institute of Food and Agricultural (NIFA; https://ror.org/03wjh4t84) Hatch Grant program (IND90024429), administered by Purdue University (https://ror.org/05p8z3f47) for partial funding support for RCW. SJB’s effort was supported by the DOE Office of Science, Office of Biological and Environmental Research (BER), Early Career Research Project grant no. SCW1883. Work at Lawrence Livermore National Laboratory was performed under U.S. Department of Energy Contract DE-AC52-07NA27344.

## DATA AVAILABILITY

The SIPdb analysis workflow (phyloseq objects and R scripts), the SQLite database and sequence data, and the dashboard app (‘app.R’) are all open source and available for local use or adapting for future uses cases via GitHub (https://github.com/alexbtrentin/sipdb-pipeline). All additional study metadata is summarized in **Table S2**, and intermediary files used for SIPdb analysis and queries, are archived on The Open Science Framework (OSF) under the accession: https://osf.io/cwg6r/overview.

