## Supplementary Methods for "SIPdb: A stable isotope probing database and analytical dashboard for linking amplicon sequences to microbial activity using a reverse ecology approach"

Trentin *et al.*, 2026

*Sequence processing and phyloseq object construction*

Raw sequencing files were retrieved from the NCBI Short Read Archive using fasterq-dump from the SRA Toolkit [1] and processed using a standardized QIIME2 workflow (v. 2024.10)[2]. Only amplicon sequences spanning the full V4 region of the 16S rRNA gene, bounded by primers 515F (5′-GTGYCAGCMGCCGCGGTAA-3′) and 806R (5′-GGACTACHVGGGTWTCTAAT-3′), were retained[3]. Quality filtering, denoising, and chimera removal were performed using the DADA2 plugin, yielding amplicon sequence variants (ASVs), and taxonomic classification was assigned against the SILVA 138.2 database[4]. Feature tables were filtered prior to differential abundance testing to remove sparse and low-abundance taxa. ASVs were retained if they met all of the following criteria: (i) at least three total counts across all samples within a study, (ii) presence in at least two samples or ≥1% of samples per study, and (iii) sparsity ≤99% across samples. Samples with fewer than 1,000 total reads were excluded together with their paired controls, and sequences annotated as chloroplasts or mitochondria were removed.

For each study, a rooted phylogenetic tree was generated using the QIIME2 q2-phylogeny workflow (align-to-tree-mafft-fasttree) to support downstream diversity analyses. Filtered ASV tables, taxonomic assignments, representative sequences, sample metadata, and phylogenetic trees were imported into R (v4.4.1)[5] and assembled into phyloseq objects (v1.48.0)[6].

*Gradient handling and window definition*

Studies were programmatically classified into one of three gradient structures based on available metadata: (case 1) binary gradients with pooled heavy and light fractions; (case 2) position-resolved gradients lacking buoyant density information; and (case 3) density-resolved gradients with measured buoyant densities. For position-resolved studies (case 2), a fixed-window approach grouped the heaviest 33% of gradient positions from labeled samples and contrasted them with the corresponding positions from unlabeled controls, excluding lighter fractions from both gradients, following Youngblut et al. (2014)[7]. Sliding-window analyses sequentially compared adjacent positions across the gradient; studies with ≤3 fractions were analyzed at the single-fraction level, whereas studies with ≥5 fractions used hierarchical clustering (Ward’s method with Euclidean distance)[8] to define two to five windows based on positional proximity.

For density-resolved studies (case 3), fixed windows were defined using buoyant density thresholds of ≥1.725 g mL⁻¹ for stringent analyses and ≥1.700 g mL⁻¹ for relaxed analyses, with optional intermediate windows (1.710–1.725 g mL⁻¹). Sliding-window analyses for density-resolved gradients used continuous density values directly and did not require clustering. An optional k-nearest-neighbor (KNN) strategy [13] selected equal numbers of labeled and unlabeled samples (default: k = 3 per group, adaptively reduced when sample numbers were limited) nearest to each gradient position.

*Statistical integration and incorporator criteria*

For each taxon and differential abundance method, effect sizes across windows were combined using inverse-variance weighting (weights = 1 / max(padj, 0.01)), and window-level p-values were combined using Fisher’s method[9]. Directional consistency was quantified as the proportion of windows showing enrichment in the same direction, and method-level evidence was integrated using Stouffer’s Z-score method[10] to generate a single cross-method consensus p-value per taxon. Taxa were designated as isotope incorporators if they met all three criteria: adjusted p-value < 0.05, absolute log₂ fold change > 1, and directional agreement across windows > 0.5. Both fixed-window and sliding-window analyses were performed for all position- and density-resolved studies, but only sliding-window consensus results were incorporated into version 1 of SIPdb; all window- and method-level outputs were retained to preserve analytical provenance.

*Validation procedures*

Synthetic microbiome datasets with defined compositions were generated and processed through the analysis pipeline. In total, 120 synthetic datasets were generated spanning three gradient configurations: binary (n = 30), position-resolved with 3–8 discrete positions (n = 45), and density-resolved gradients (n = 45). Synthetic communities varied in richness (100–220 ASVs per dataset), sample number (16–48 samples), and isotope incorporation patterns. For each dataset, 5–18% of ASVs were designated as true incorporators, with simulated enrichment magnitudes of 2.8–5.5-fold in labeled samples from heavy fractions (buoyant density ≥1.725 g mL⁻¹ or gradient positions 1–2). Biological noise was introduced using negative binomial count distributions with dispersion parameters typical of amplicon sequencing data (size = 10), and natural abundance control samples were simulated without enrichment. Each synthetic dataset was processed through the full analysis pipeline, and outputs were compared against known ground-truth incorporator identities to calculate sensitivity, precision, specificity, F1 score, and Matthews correlation coefficient [9].

Pipeline outputs were compared to isotope incorporators reported in the original publications describing each included dataset. For each study, manuscripts and supplementary materials were systematically reviewed to extract reported isotope-incorporating taxa. When available, ASV-level identifiers were retrieved directly; otherwise, incorporators were recorded at the genus or family level. Reported taxa were queried against SIPdb to determine whether they were detected as significant incorporators (padj ≤ 0.05, log₂FC > 1) in the corresponding study and isotopolog using any of the four differential abundance methods implemented in the pipeline. A reported taxon was considered validated if it appeared in the SIPdb results for the matching study and isotopolog (Supplementary Table S3), and the detecting methods and mean log₂FC value were recorded.

*Database construction and access*

Differential abundance results generated for each study, window, and analytical method were compiled into a relational SQLite database with a normalized schema. All analyses were performed at the ASV level, with taxonomic incorporator designations derived from ASV results. Sliding-window analyses using the k-nearest-neighbor (KNN) strategy selected k = 3 samples per group nearest to each gradient position. Window metadata included position ranges, window mode, and sample sizes per group. Studies were excluded prior to database ingestion if they had insufficient replication (<2 samples per condition), extreme sparsity (>98% zeros), or lacked appropriate unlabeled controls.

For each study, quality control metrics were recorded, including the number of labeled and unlabeled samples, median library size, data sparsity, and the number of ASVs passing filtering thresholds. The primary results tables store differential abundance statistics linked by foreign keys to study metadata, taxonomic annotations, and representative sequences. Representative V4 region sequences were compiled into a BLAST nucleotide database, and user-supplied sequences were matched using BLASTn with identity thresholds of ≥97% for genus-level matches or 100% identity across the full V4 region for exact ASV-level matches. Prior to finalizing SIPdb version 1, database integrity checks confirmed output completeness, study coverage, and file artifact integrity.

*Dashboard implementation and downstream analyses*

The SIPdb analysis dashboard was implemented as a Shiny web application [11] backed by a SQLite database. The application interfaces directly with curated SIPdb tables and supports interactive filtering based on statistical thresholds, density threshold, and differential abundance method. Tabular outputs and visual summaries are generated using ggplot2 [12], plotly [13], and viridisLite [14], with all results exportable in CSV format. Downstream analyses using curated SIPdb outputs were performed as described in the Main Methods. Principal component analysis was conducted on scaled and centered variables using the prcomp function in R following log₁₀ transformation of count-based variables. Non-metric multidimensional scaling (NMDS) was performed on phylum-level incorporator relative abundances using Bray–Curtis dissimilarity with two dimensions and 100 random starts in the vegan package (v2.6-4) [15], and community differences were tested using PERMANOVA with 999 permutations. Re-analysis of non-SIP datasets followed the same processing and annotation procedures described above [16].
